# Crystal structures of SARS-CoV-2 ADP-ribose phosphatase (ADRP): from the apo form to ligand complexes

**DOI:** 10.1101/2020.05.14.096081

**Authors:** Karolina Michalska, Youngchang Kim, Robert Jedrzejczak, Natalia I. Maltseva, Lucy Stols, Michael Endres, Andrzej Joachimiak

**Author notes:** These authors provided equal contribution (Karolina Michalska, Youngchang Kim). Correspondence should be addressed to: Andrzej Joachimiak, Structural Biology Center, X-ray Science Division Argonne National Laboratory, Argonne, Illinois 60439, USA.

## Abstract

Among 15 nonstructural proteins (Nsps), the newly emerging SARS-CoV-2 encodes a large, multidomain Nsp3. One of its units is ADP-ribose phosphatase domain (ADRP, also known as macrodomain) which is believed to interfere with the host immune response. Such a function appears to be linked to the protein’s ability to remove ADP-ribose from ADP-ribosylated proteins and RNA, yet the precise role and molecular targets of the enzyme remains unknown. Here, we have determined five, high resolution (1.07 - 2.01 Å) crystal structures corresponding to the apo form of the protein and complexes with 2-(N-morpholino)ethanesulfonic acid (MES), AMP and ADPr. We show that the protein undergoes conformational changes to adapt to the ligand in a manner previously observed before for in close homologs from other viruses. We also identify a conserved water molecule that may participate in hydrolysis. This work builds foundations for future structure-based research of the ADRP, including search for potential antiviral therapeutics.

## Introduction

Over the past several months, Severe Acute Respiratory Syndrome Coronavirus 2 (SARS-CoV-2) has been spreading across the world causing a disease termed COVID-19 (Coronaviridae Study Group of the International Committee on Taxonomy of Viruses, 2020). The virus emerged first in December of 2019 in Wuhan, China, and since then, has rapidly traveled to most countries around the world, infecting millions and killing hundreds of thousands. These developments forced The World Health Organization to declare the outbreak as a pandemic. In the absence of natural community immunity, tested vaccine or approved drugs that would help to control the epidemics, billions of people are currently under quarantine or lockdowns to minimize further transmissions.

The etiological agent of COVID-19 was isolated and identified as a novel coronavirus resembling SARS-CoV responsible for the 2002-2003 outbreak (Wu *et al.*, 2020). As other coronaviruses, SARS-CoV-2 utilizes positive-sense RNA genome encoding nonstructural proteins (Nsps), structural proteins, such as spike glycoprotein (S), envelope (E), membrane (M) and nucleocapsid proteins (N), as well accessory proteins (Wu *et al.*, 2020). Nsps are encompassed within ORF1a and ORF1b, that produce two polyproteins, pp1a and pp1ab (Cui *et al.*, 2019). The latter protein results from the ribosomal shift that enables continuous translation of ORF1a along with ORF1b. Pp1a contains two viral proteases, 3C-like main protease (Mpro, encoded by Nsp5), and papain-like protease (PLpro, encoded within Nsp3), responsible for posttranslational processing of two polyproteins. The cleavage yields 16 Nsps (15 Nsps in SARS-CoV-2) (Baez-Santos *et al.*, 2015) that form a large membrane-bound replicase complex.

The largest component of the replicase assembly is Nsp3. This multidomain protein, among other modules (N-terminal Nsp3a domain, ADP-ribose phosphatase domain (ADRP, also known as macrodomain), SARS-unique domain, PLpro, RNA binding domain, marker domain, transmembrane domain and Y-domain (https://coronavirus3d.org). ADRP was identified thirty years ago by bioinformatics as a unique and conserved domain, initially termed X domain (Lee *et al.*, 1991), found in the genomes of *Togaviridae, Coronaviridae*, and *Hepeviridae* families. Since then it has been also discovered in *Iridiviridae, Poxviridae* and *Myoviridae*, which includes phages. Crystallographic model of the first ADRP (Saikatendu *et al.*, 2005) enabled to the classification of the protein as a member of macroH2A-like family. The founding member of the family is a large, nonhistone part of the histone macroH2A, known as macrodomain (Pehrson & Fried, 1992). Until now, structurally characterized viral ADRPs cover 11 representatives, including those from SARS-CoV, MERS-CoV (Middle East Respiratory Syndrome virus), H-CoV-229E (Human Coronavirus 229E), and others.

The non-viral macrodomains have been shown to recognize ADP-ribose (ADPr) in a free form and in macromolecule-linked forms, as well as attached to other ligands. Besides binding, some macrodomains possess catalytic activities, including the removal of ADPr from ADP-ribosylated proteins or nucleic acids (DNA and RNA). ADP-ribosylation is a regulatory modification present in all kingdoms of life and is known to play a role in the DNA damage repair, signal transduction, immune response and other cellular stresses (for review, see for example (Crawford *et al.*, 2018)). The appendage is transferred onto the target by ADP-ribosyl transferases (ARTs) classified as either diphtheria toxin-like (ARTD, previously known as poly(ADP-ribose)polymerases, PARPs) or cholera toxin-like enzymes (ARTC). Some sirtuins also carry out such reactions. Both groups of proteins utilize NAD^+^ as an ADPr donor. ARTD catalyze the transfer of either single (mono-ADP-ribosylation, MARylation) or multiple ADPr units (poly-ADP-ribosylation, PARylation, mostly PARP1 and PARP2), primarily onto glutamate/aspartate residues, but sometimes also serine. In nucleic acids, the modification is attached to phosphate group at the terminal ends of DNA/RNA. ARTCs carry out only MARylation and preferentially act on arginine. The de-ADP-ribosylation requires several enzymes. The polymeric fragment of the modification is removed by poly(ADP-ribosyl)glycohydrolase (PARG) while the final ADP-ribose unit is cut off from glutamate/aspartate residues by macrodomains. Enzymes from family of (ADP-ribosyl) hydrolases (ARHs) have specificity for serine and arginine cargos (Fontana *et al.*, 2017, Moss *et al.*, 1988). The released ADPr is used in recycling pathways.

Currently, six macrodomains classes are distinguished: MacroH2-like, AlC1-like, PARG-like, Macro2-type, SUD-M-like (also known as Mac2/Mac3), and MacroD-type (Rack *et al.*, 2016). These categories are derived from structural similarities rather than sequence similarity. Most viral macrodomains fall into the MacroD-like family, encompassing human homologs MacroD1 and MacroD2, associated with the removal of mono(ADP-ribosylation). *In vivo* experiments showed that viral MacroD-like macrodomains can hydrolyze ADPr-1”-phosphate, yet the catalytic efficiency of this process raised doubts about its physiological implications (Egloff *et al.*, 2006). Instead, it has been suggested that these macrodomains might play roles analogous to MacroD1 and MacroD2. Indeed, de-ADP-ribosylating activities on protein and RNA, including the removal of the entire PAR chain, have been demonstrated for several viral macrodomains, for example those from SARS-CoV and H-CoV 229E (Li *et al.*, 2016, Eckei *et al.*, 2017, Munnur *et al.*, 2019). Binding to PAR (Egloff *et al.*, 2006) and RNA (Malet *et al.*, 2009) has been also reported. Importantly, a wide range of affinities and activities practically prevents similarity-based assumptions about physiological role of these proteins.

Such biochemical activity, on physiological level means that the role of ADRP would be to counteract function of the ARTD/PARP proteins. The latter enzymes are upregulated by interferon, indicating their relevance in innate immune response. The systematic knockdown studies of all 17 mammalian PARP in the MHV model with macrodomain mutants implicated PARP12 and PARP14 in the control of virus replication (Grunewald *et al.*, 2019). Interestingly, PARP12, which is able to auto-ADP-ribosylate, belongs to a family of zinc finger CCCH domain known to bind to RNAs, including those of viral origin. The antiviral properties of the protein have been linked to its enzymatic activity, colocalization with polyribosomes via RNA-binding domain and interference with translation machinery (Atasheva *et al.*, 2014). Also, PARP10, known to modify RNA (Munnur *et al.*, 2019), has been shown to inhibit viral replication (Atasheva *et al.*, 2012, Atasheva *et al.*, 2014). The role of ADRP in jeopardizing immune response has also been emphasized by studies showing that viruses with mutated macrodomain replicated poorly in bone-marrow-derived macrophages, which are the primary cells mounting an innate immune response (Grunewald *et al.*, 2019). Along the same lines, viruses with deactivated macrodomains were sensitive to interferon (IFN) pretreatment (Kuri *et al.*, 2011).

Since the role of macrodomains in pathogenesis is essential, it appears that their inhibition may help to reduce viral load and facilitate recovery. Therefore, these proteins might be attractive targets for the developments of small molecule antivirals, assuming that highly selective compounds could be found discriminating between viral and human macrodomains. As a step towards this goal, we have determined crystal structure of SARS-CoV-2 ADRP in multiple states: in the apo form, complex with 2-(N-morpholino)ethanesulfonic acid (MES), AMP and ADPr. With the apo crystals diffracting to atomic resolution, we have developed a robust system for structure-based experiments to identify potential small molecule inhibitors.

## Results and Discussion

### Protein production and structure determination

We have used *E. coli* codon-optimized synthetic gene with a sequence corresponding to SARS-CoV-2 ADRP to produce protein for crystallographic and biochemical studies. The protein has been crystallized under several conditions, yielding five crystal structures, denoted as ADRP/APO1 (apo form), ADRP/ADPr (complex with ADPr), ADRP/AMP (complex with AMP), ADRP/MES (complex with 2-(*N*-morpholino)ethanesulfonic acid) and ADRP/APO2 (apo form). The ADRP/APO1 structure was solved first, by molecular replacement with the SARS-CoV homolog structure (PDB id 2ACF) as a search model. All the subsequent structures were solved by MR with refined SARS-CoV-2 ADRP structure as a template.

ADRP/APO1 has been refined to 2.01 Å resolution. The protein crystallized in *P*1 space group with two molecules in the unit cell. None of the polypeptides contains ligand in the catalytic pocket, but there is a *N*-cyclohexyl-2-aminoethanesulfonic acid (CHES) molecule bound on the surface. As ADRP/APO1, the ADRP/ADPr structure was solved in *P*1 space group. It was refined with reflections extending to 1.50 Å, though 88% completeness is achieved only to 1.65 Å resolution. In both polypeptide chains, the ADPr ligand is well-defined in the electron density map. ADRP/AMP crystallized in *P*21 space group also with two molecules in the asymmetric unit. The atomic model was refined to 1.45 Å. In the ADPr-binding pocket, one of the protein molecules (chain A) binds AMP ligand with 0.8 occupancy, while the other (chain B) binds MES molecule with occupancy 0.7. In the latter case, there is an additional electron density in the position where adenine ring binds, but its quality prevented acceptable interpretation. The ADRP/MES crystals also grew in *P*21, but with smaller unit cell and only one protein molecule in the asymmetric unit. These crystals diffracted to 1.07 Å resolution. In the structure two MES molecules have been identified: one in the ADPr-binding pocket and another on the protein surface. Finally, the ADRP/APO2 structure has been determined in C2 space group with one protein chain in the asymmetric unit. In the refinement we used reflections extending to 1.35 Å resolution. In ADRP/APO2, the binding pocket has no small molecule present, with exception of solvent. In all structures, the polypeptide chains are nearly complete, with only a few residues missing at the termini, as detailed in the Methods section. The data collection and structure refinement statistics are given in Table 1. All the structures have been deposited in the Protein Data Bank (PDB).

**Table 1.**
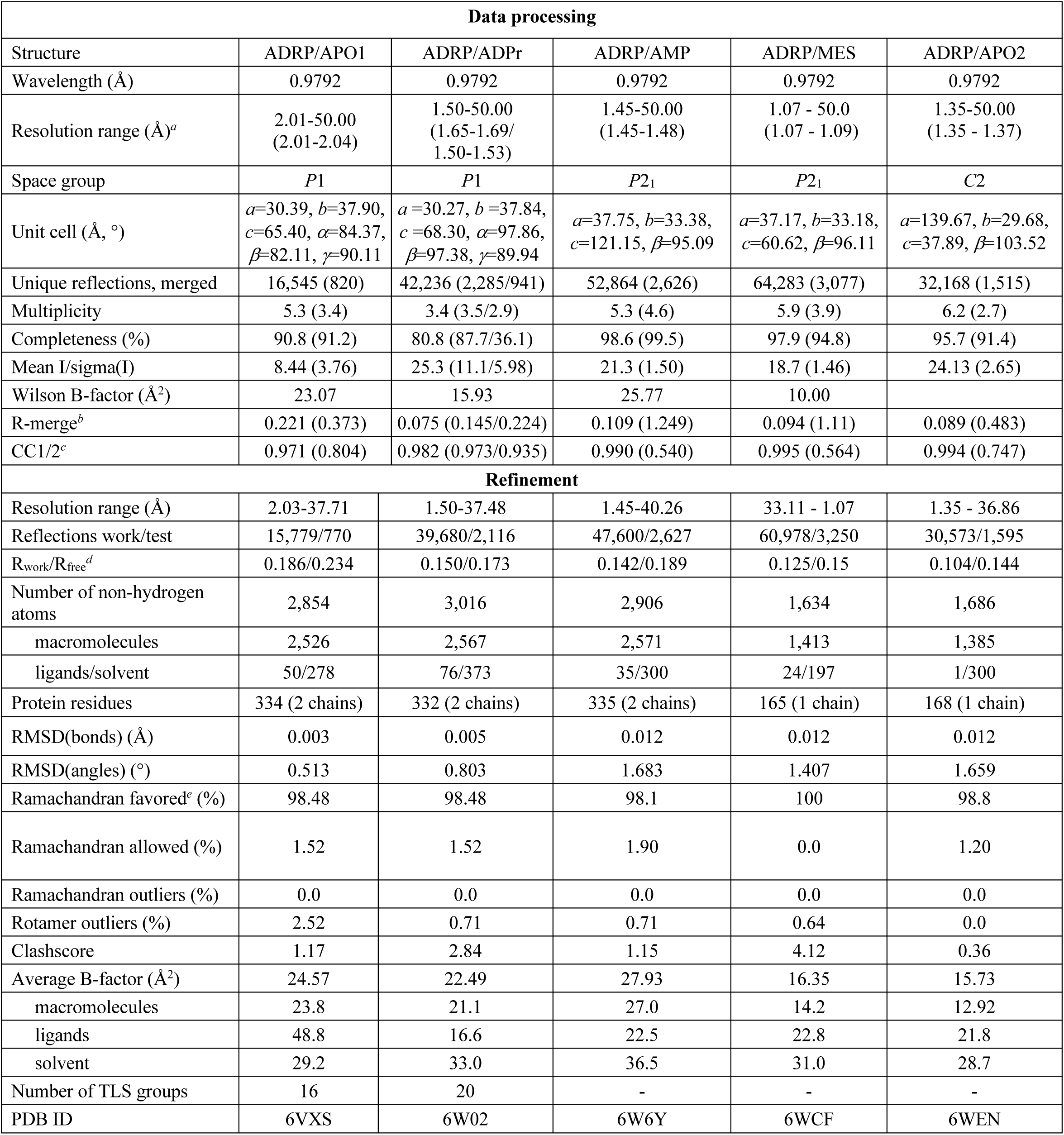
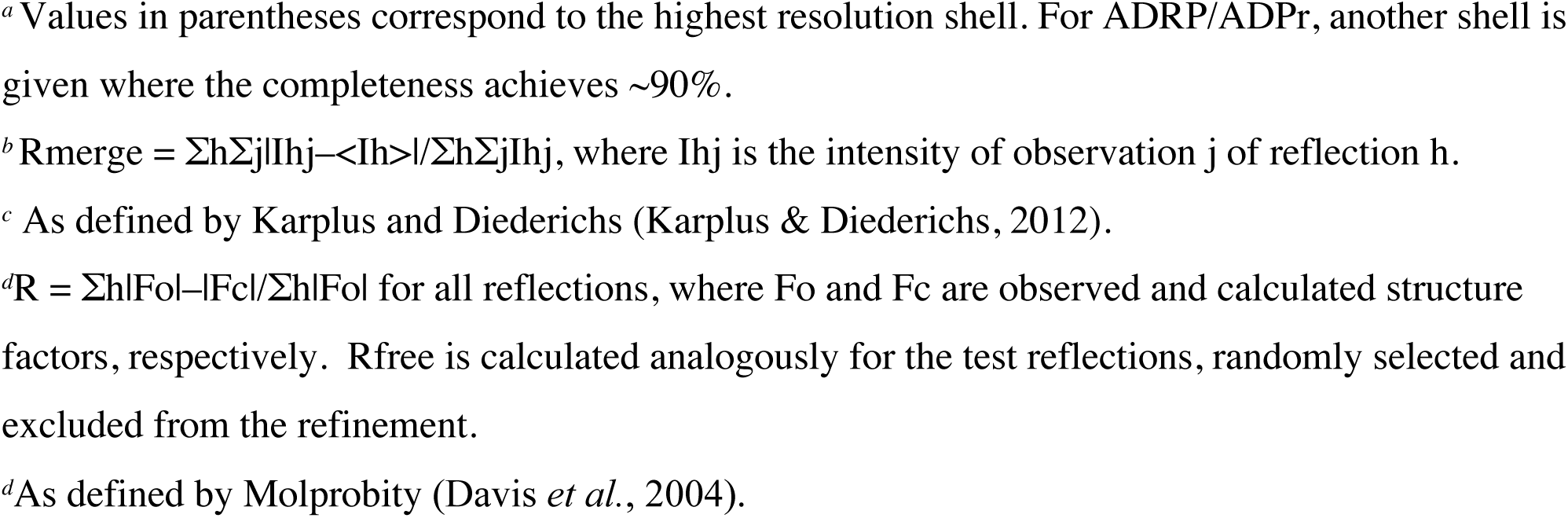
Data processing and refinement statistics.

| Data processing |  |  |  |  |  |
| --- | --- | --- | --- | --- | --- |
| Structure | ADRP/APO1 | ADRP/ADPr | ADRP/AMP | ADRP/MES | ADRP/APO2 |
| Wavelength (Å) | 0.9792 | 0.9792 | 0.9792 | 0.9792 | 0.9792 |
| Resolution range (Å)<sup>a</sup> | 2.01-50.00<br>(2.01-2.04) | 1.50-50.00<br>(1.65-1.69/<br>1.50-1.53) | 1.45-50.00<br>(1.45-1.48) | 1.07 - 50.0<br>(1.07 - 1.09) | 1.35-50.00<br>(1.35 - 1.37) |
| Space group | *P*1 | *P*1 | *P*2<sub>1</sub> | *P*2<sub>1</sub> | *C*2 |
| Unit cell (Å, °) | *a*=30.39, *b*=37.90,<br>*c*=65.40, $\alpha$=84.37,<br>$\beta$=82.11, $\gamma$=90.11 | *a* =30.27, *b* =37.84,<br>*c* =68.30, $\alpha$=97.86,<br>$\beta$=97.38, $\gamma$=89.94 | *a*=37.75, *b*=33.38,<br>*c*=121.15, $\beta$=95.09 | *a*=37.17, *b*=33.18,<br>*c*=60.62, $\beta$=96.11 | *a*=139.67, *b*=29.68,<br>*c*=37.89, $\beta$=103.52 |
| Unique reflections, merged | 16,545 (820) | 42,236 (2,285/941) | 52,864 (2,626) | 64,283 (3,077) | 32,168 (1,515) |
| Multiplicity | 5.3 (3.4) | 3.4 (3.5/2.9) | 5.3 (4.6) | 5.9 (3.9) | 6.2 (2.7) |
| Completeness (%) | 90.8 (91.2) | 80.8 (87.7/36.1) | 98.6 (99.5) | 97.9 (94.8) | 95.7 (91.4) |
| Mean I/sigma(I) | 8.44 (3.76) | 25.3 (11.1/5.98) | 21.3 (1.50) | 18.7 (1.46) | 24.13 (2.65) |
| Wilson B-factor (Å<sup>2</sup>) | 23.07 | 15.93 | 25.77 | 10.00 |  |
| R-merge<sup>b</sup> | 0.221 (0.373) | 0.075 (0.145/0.224) | 0.109 (1.249) | 0.094 (1.11) | 0.089 (0.483) |
| CC1/2<sup>c</sup> | 0.971 (0.804) | 0.982 (0.973/0.935) | 0.990 (0.540) | 0.995 (0.564) | 0.994 (0.747) |
| Refinement |  |  |  |  |  |
| Resolution range (Å) | 2.03-37.71 | 1.50-37.48 | 1.45-40.26 | 33.11 - 1.07 | 1.35 - 36.86 |
| Reflections work/test | 15,779/770 | 39,680/2,116 | 47,600/2,627 | 60,978/3,250 | 30,573/1,595 |
| R<sub>work</sub>/R<sub>free</sub><sup>d</sup> | 0.186/0.234 | 0.150/0.173 | 0.142/0.189 | 0.125/0.15 | 0.104/0.144 |
| Number of non-hydrogen atoms | 2,854 | 3,016 | 2,906 | 1,634 | 1,686 |
| macromolecules | 2,526 | 2,567 | 2,571 | 1,413 | 1,385 |
| ligands/solvent | 50/278 | 76/373 | 35/300 | 24/197 | 1/300 |
| Protein residues | 334 (2 chains) | 332 (2 chains) | 335 (2 chains) | 165 (1 chain) | 168 (1 chain) |
| RMSD(bonds) (Å) | 0.003 | 0.005 | 0.012 | 0.012 | 0.012 |
| RMSD(angles) (°) | 0.513 | 0.803 | 1.683 | 1.407 | 1.659 |
| Ramachandran favored<sup>e</sup> (%) | 98.48 | 98.48 | 98.1 | 100 | 98.8 |
| Ramachandran allowed (%) | 1.52 | 1.52 | 1.90 | 0.0 | 1.20 |
| Ramachandran outliers (%) | 0.0 | 0.0 | 0.0 | 0.0 | 0.0 |
| Rotamer outliers (%) | 2.52 | 0.71 | 0.71 | 0.64 | 0.0 |
| Clashscore | 1.17 | 2.84 | 1.15 | 4.12 | 0.36 |
| Average B-factor (Å<sup>2</sup>) | 24.57 | 22.49 | 27.93 | 16.35 | 15.73 |
| macromolecules | 23.8 | 21.1 | 27.0 | 14.2 | 12.92 |
| ligands | 48.8 | 16.6 | 22.5 | 22.8 | 21.8 |
| solvent | 29.2 | 33.0 | 36.5 | 31.0 | 28.7 |
| Number of TLS groups | 16 | 20 | - | - | - |
| PDB ID | 6VXS | 6W02 | 6W6Y | 6WCF | 6WEN |

<sup>a</sup> Values in parentheses correspond to the highest resolution shell. For ADRP/ADPr, another shell is given where the completeness achieves ~90%.
<sup>b</sup> $R_{\text{merge}} = \sum_h \sum_j |I_h - \langle I_h \rangle| / \sum_h \sum_j I_h$ , where $I_h$ is the intensity of observation $j$ of reflection $h$ .
<sup>c</sup> As defined by Karplus and Diederichs (Karplus & Diederichs, 2012).
<sup>d</sup> $R = \sum_h |F_o - F_c| / \sum_h F_o$ for all reflections, where $F_o$ and $F_c$ are observed and calculated structure factors, respectively. $R_{\text{free}}$ is calculated analogously for the test reflections, randomly selected and excluded from the refinement.
<sup>d</sup> As defined by Molprobit (Davis *et al.*, 2004).

### Overall structure

The structure of SARS-CoV-2 ADRP features a central 7-stranded mixed β-sheet (β1↑, β2↓, β7↓, β6↓, β3↓, β5↓, β4↑) sandwiched between two layers of helices: α1, α2 and α3 on one side and η1, α4/η2, η3, α5 and α6 on the other (Fig. 1). These features follow previously established characteristic fold of a MacroD-like macrodomain, described earlier for several viral homologs. According to DALI calculations (Holm & Rosenstrom, 2010), the closest structural relative comes from SARS-CoV (2ACF, (Saikatendu *et al.*, 2005), Z-score 33.9, rmsd 0.5 Å over 168 Cα atoms superposed onto ADRP/APO2). This homolog shares 71% sequence identity (82% similarity) with the SARS-CoV-2 ADRP (as determined by EMBOSS Needle (Rice *et al.*, 2000)). Next hit corresponds to the MERS-CoV homolog (5HIH, Z-score 28.0, rmsd 1.3 Å over 163 Cα atoms, (Lei & Hilgenfeld, 2016)), that displays 40% sequence identity (61% similarity). Subsequent neighbors with rmsd up to 2 Å include *Tylonycteris* bat coronavirus HKU4 (6MEN, DOI: 10.2210/pdb6MEN/pdb), feline coronavirus (FIP, 3EW5, (Wojdyla *et al.*, 2009)), H-CoV-229E (3EJG, (Piotrowski *et al.*, 2009)), and H-CoV-NL63 (2VRI, 10.2210/pdb2VRI/pdb).

**Fig. 1.**
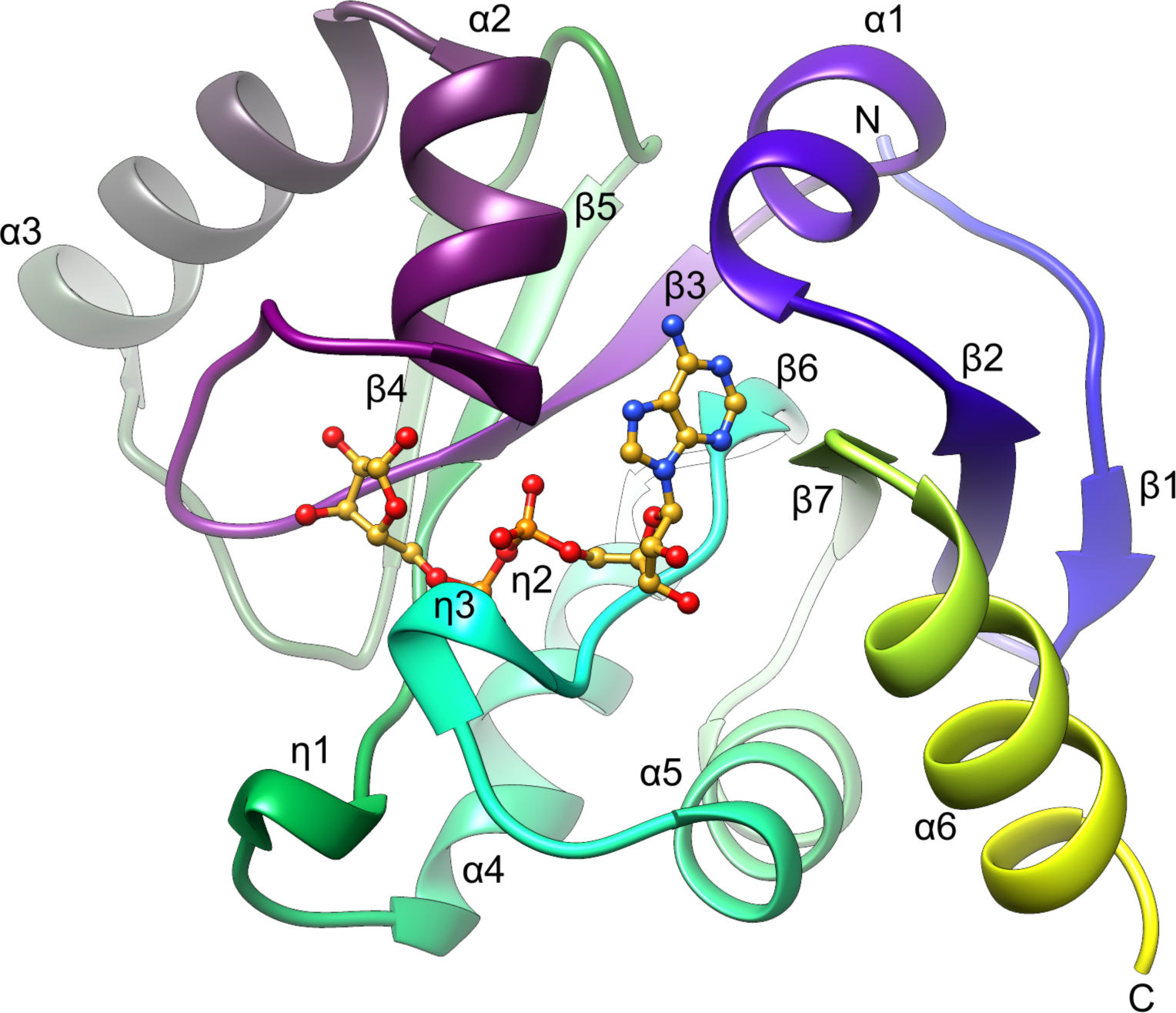
The structure of SARS-CoV-2 ADRP. The ribbon diagram shows ADRP/APO2. The ADPr ligand molecule is shown based on the superposition with the ADRP/ADPr structure.

The SARS-CoV-2 ADRP structures show high level of agreement between each other. Rmsds for ADRP/APO2 superpositions range from 0.3 and 0.4 Å (ADRP/AMP), through 0.4 and 0.5 Å (ADRP/ADPr, ADRP/MES) up to 0.6 and 0.7 Å (ADRP/APO1).

### Substrate binding pocket

The well-defined substrate binding pocket is created by C-terminal edges of the central β strands β3, β5, β6, β7, and surrounding fragments, primarily loop β3-α2, N-terminus of α1 and a long loop connecting β6 with α5, which encompasses short 310 helix η3. These elements encompass four conserved sequence motifs (Fig. 2) shared by the family members (Saikatendu *et al.*, 2005). First such block is present at the end of β3 and is followed by another, extending onto the helix α2. Third segment corresponds to the end of β5 and the last one overlaps with helix η3.

Within the crevice, four sections can be distinguished, corresponding to adenine-, distal ribose-, diphosphate- and proximal ribose-binding sites, denoted here as A, R1, P1-P2, R2. The ADRP/ADPr structure illustrates how the ligand molecule interacts with these subsites (Fig. 3, Fig. 4). The adenine moiety is sandwiched between α2 and β7 in a mostly hydrophobic environment created by I23, V49, P125, V155 and F156. Polar contacts are facilitated by D22, which forms a hydrogen bond with N6 atom via its carboxylate group and by main chain amide of I23 which binds to N1. In addition, water-mediated contacts link N3 with main chain of A154 and L126. The A site has limited sequence conservation: only P125 and D22 are conserved among homologs. Other hydrophobic residues are replaced by side chains with similar chemical character. The striking exception is F156, which in the closest homologs, SARS-CoV and MERS-CoV, is replaced by N. In other virial representatives it is substituted by another hydrophobic residue. The distal ribose ring participates only in water mediated hydrogen bonds with main chain amide of L126 and carbonyl group of A154 via the ring oxygen atom, and D157 main chain and side chain via the OH2’ group. The diphosphate moiety binds between two loops, β3-α2 and β6-(η3)-α5, that cover three segments with high sequence conservation, including glycine-rich segment (Gly46-Gly47-Gly48) within the former loop. Here, the ligand forms direct hydrogen bonds to the main chain amides of V49, S128, G130, I131, P132, and water-mediated contacts with A38, A39, A50, V95, G97. An elaborate network of water molecules also links diphosphate to G47, A129 and D157. Finally, the proximal ribose ring is stabilized in the pocket by hydrophobic interactions with F132 and I131 as well as a set of hydrogen bonds with G46 (OH2’), G48 (OH1’), N40 (OH3’). All of these residues are conserved. Additional bonds to main chain peptides of N40, K44, A50 are water-mediated. Interestingly, as described above, only few hydrogen bonds involve protein side chains, most of such contacts utilize main chain atoms. This may explain why there is less pressure on amino acid sequence preservation, since main chain interactions can be accomplished with multiple side chains combinations.

**Fig. 2.**
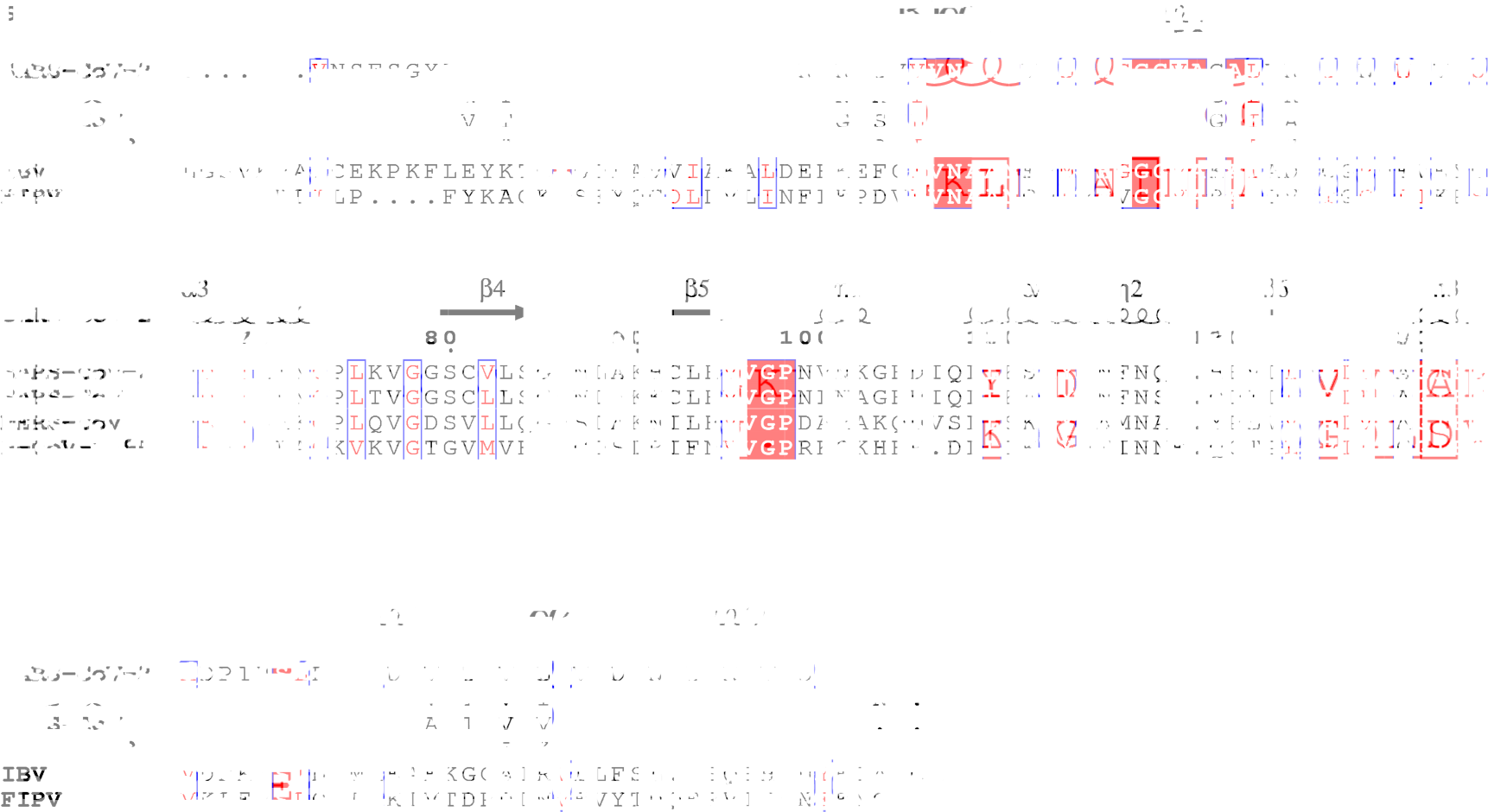
Sequence alignment of SARS-CoV-2 ADRP homologs from coronaviruses with structures in complexes with ADPr available in the PDB: SARS-CoV-2 (6WEN, chain A), SARS-CoV (2AFAV, chain A), MERS-CoV (5DUS, chain A), H-CoV-229E (3EWR, chain A), IBV (3EWP, chain A) FIPV (3JZT, chain A). The secondary structure elements labeled for SARS-CoV-2 ADRP.

**Fig. 3.**
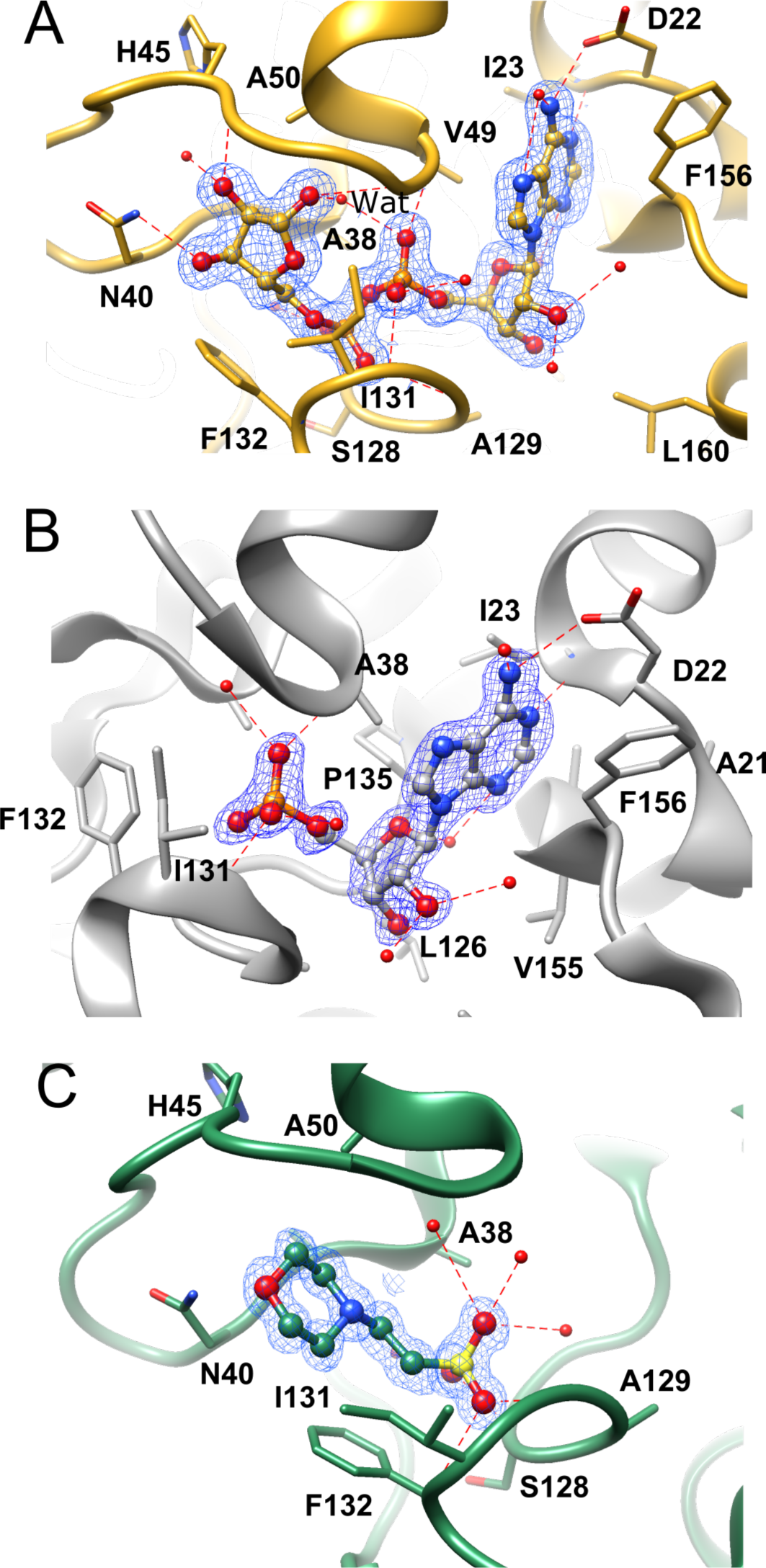
Ligand binding in the SARS-CoV-2 ADRP. A). ADPr binding (chain A, 6W02). B). AMP binding (chain A, 6W6Y). C). MES binding (6WCF). All 2mFo-DFc electron density maps are contoured at 1.2 σ level.

**Fig. 4.**
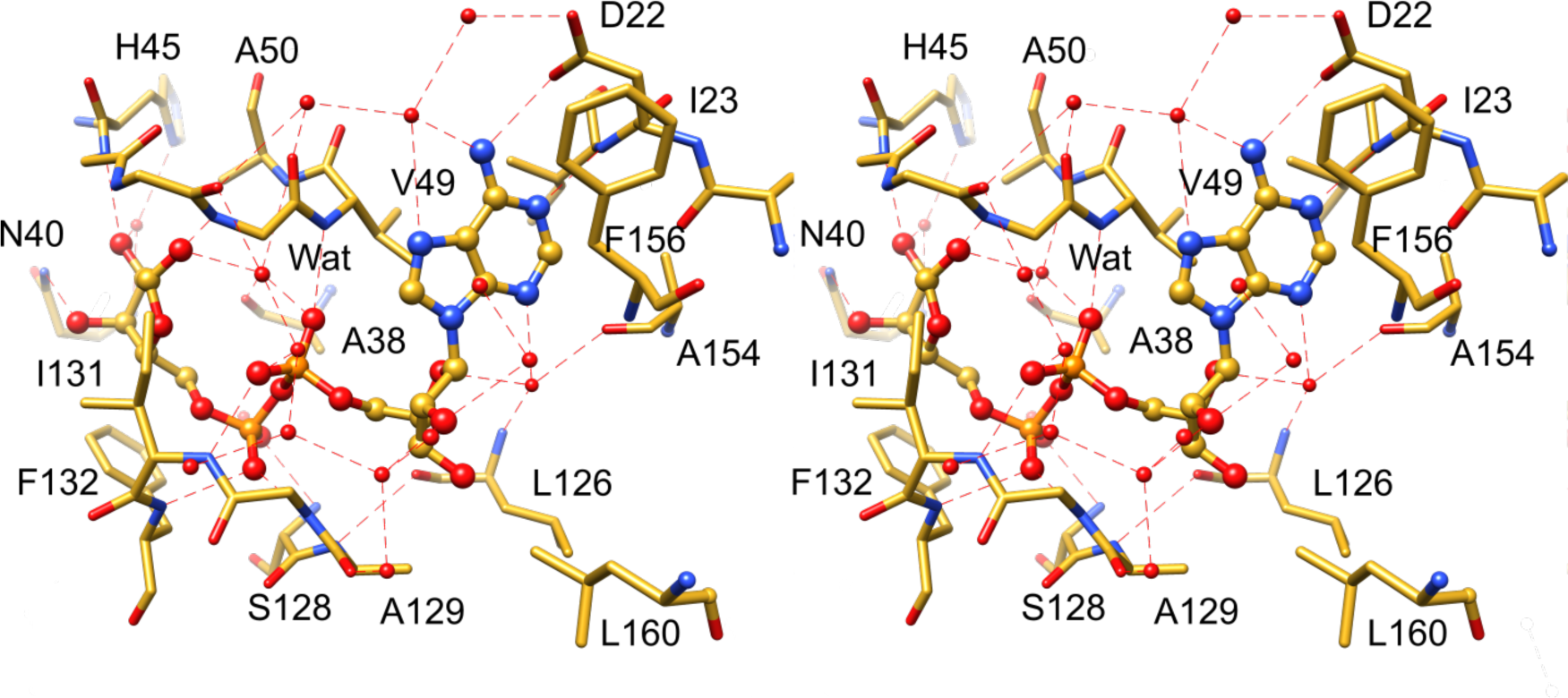
Stereoview of the ADPr binding in the SARS-CoV-2 ADRP binding site.

Similar contacts are observed in the ADRP/AMP structure (Fig. 3), in which the ligand superposes well with the AMP portion of the ADPr ligand (Fig. 5). The ADRP/MES complex though, presents somewhat different scenario, where the 2-*N*-morpholine ring takes place of the proximal ribose and sulfonic acid substitutes distal phosphate. The latter group forms hydrogen bonds observed in the ADPr complex, and an additional network of solvent-facilitated contacts. The ring moiety appears to primarily be anchored by hydrophobic interactions with F132 and I131, potentially a hydrogen bond might be present between the morpholine oxygen atom and N40, though geometry is rather unfavorable.

**Fig. 5.**
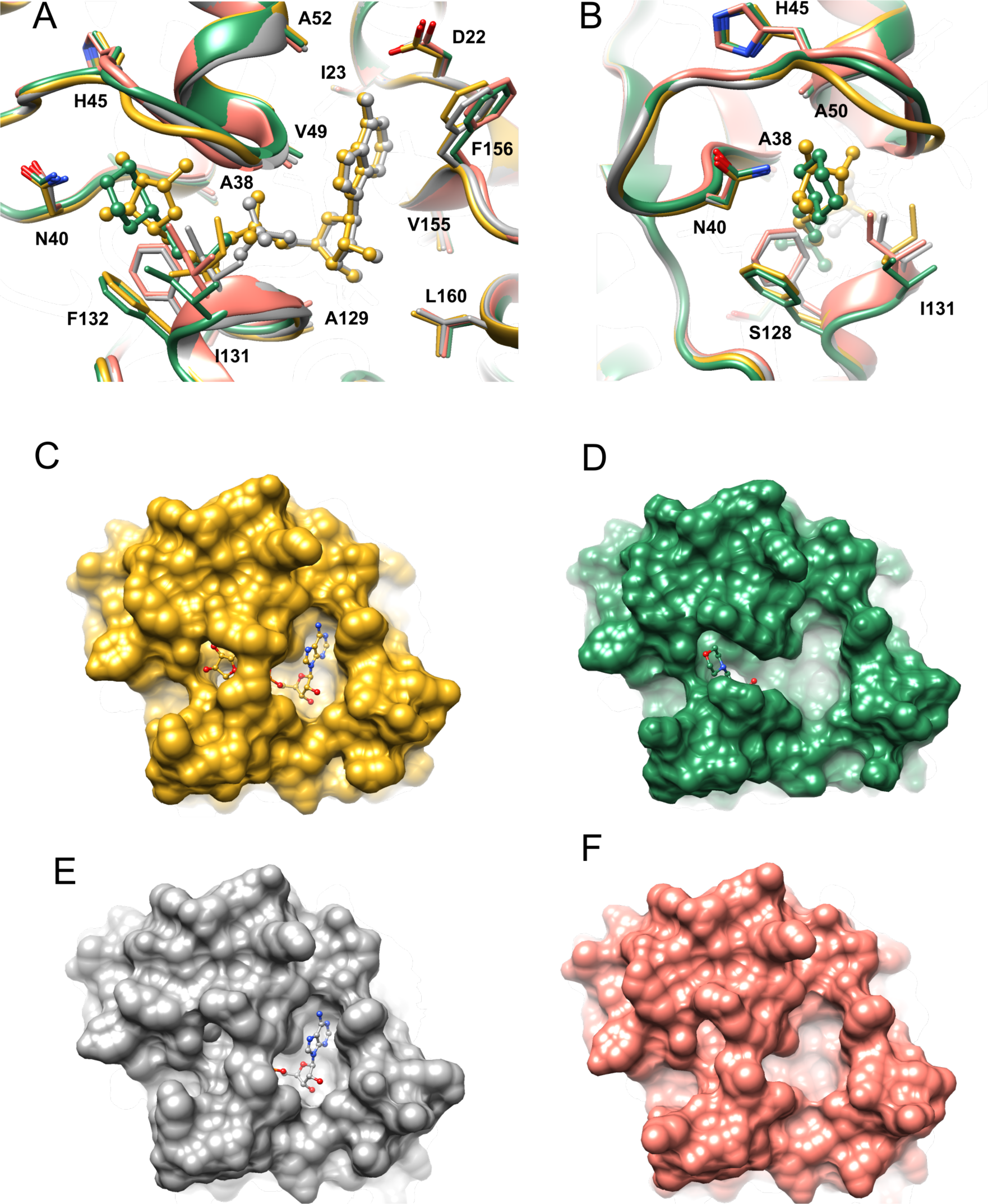
Ligand-induced conformational changes in the SARS-CoV-2 ADRP structure. **A).** Superposition of ADRP/ADPr complex (yellow, chain A, 6W02) with ADRP/AMP (grey, chain A, 6W6Y), ADRP/MES (green, 6WCF) and ADRP/APO2 (coral, 6WEN). The ligand molecules are shown in ball-and-stick representation. B). As in A, but rotated ∼90°. C). Surface represenation of ADRP/ADPr complex. D). Surface represenation of ADRP/MES complex. E). Surface represenation of ADRP/AMP complex. F). Surface represenation of ADRP/APO2 structure.

### Ligand-induced conformational changes

While interactions with ligands do not trigger major conformational changes in the overall structure, significant shifts are observed in the binding pocket itself. Superpositions of the apo forms with the complexed proteins indicate several adjustments (Fig. 5). First, in the A site, F156 is brought closer to the pocket lumen when it is occupied by the nucleotide, as seen in ADRP/ADPr and ADRP/AMP complexes. Then, the glycine-rich β3-α2 loop shows a high degree of flexibility with roughly the same geometry but slightly different positions in ADRP/APO1, ADRP/MES, ADRP/AMP, and ADRP/APO2 (Fig. 5). In the latter structure, though, the Gly46-Gly47 peptide bond has also an alternative conformation. Significant change is observed in ADRP/ADPr, where the loop has to rearrange to make main-chain amide nitrogen atoms accessible for interactions with ribose OH1’ and OH2’ groups. Finally, the geometry of β6-(η3)-α5 loops and rotameric states of F132 and I131, contributing to P1-P2 and R2 sites, also adapts depending on the ligand identity. The apo and AMP-bound forms contain the η3 element within the β6-α5 linker, while in the ADPr and MES complexes this region does not observe helix 310 parameters. The primary reason for this is the flipping A129-G130 peptide bond, which in the absence of phosphate 2, or its mimetic, has the carbonyl group facing P2 site. Otherwise, with P2 occupied, the G130 amide group is hydrogen-bonded with the ligand, as described above. I131 and F132 are also observed in two states. With the R2 pocket empty or with MES, I131 adopts *pt* rotamer (*p*: plus, centered near +60°; *t*: trans, centered near 180°), while in the presence of a ribose ring, it turns into the *mt* state (*m*: minus, centered near −60°) (Hintze *et al.*, 2016). F132 follows somewhat similar pattern: in the first scenario is adopts *m*-10 conformation, while in the latter, *m*-80. These rearrangements are necessary to provide sufficient room for the ligand and proper interactions. Similar transformations in the ligand-binding pocket have been reported for other homologs (Egloff *et al.*, 2006, Piotrowski *et al.*, 2009, Wojdyla *et al.*, 2009). In the ADRP/APO1 structure, while the described geometry of the β6-η3-α5 linker remains similar to ADRP/APO2, the entire section and neighboring η1 are shifted away from the binding pocket.

### Similarity of ADPr binding between ADRP homologs

The PDB currently holds four other coronaviral ADRPs in complexes with ADPr: from SARS-CoV (2FAV, (Egloff *et al.*, 2006)), MERS-CoV (5HOL, (Lei *et al.*, 2018), 5DUS, (Cho *et al.*, 2016)), H-CoV-229E (3EWR, (Xu *et al.*, 2009)), and also animal infecting IBV (3EWP, (Piotrowski *et al.*, 2009)) and FIPV (3JZT, (Wojdyla *et al.*, 2009)). The SARS- and MERS-CoV complexes mostly follow the pattern of interactions observed in the current structure (Fig. 6). The ligand geometry is also preserved. The elements that are distinct are located in the A and R1 sites. Most strikingly, SARS-CoV-2 F156 is in the SARS-CoV homolog replaced by N157 (N154 in MERS-CoV 5DUS numbering) that stacks against the adenine ring and at the same time created water-mediated hydrogen-bonds with the distal ribose. In that region, three other sequence discrepancies with the MERS-CoV ADRP are located: I23 is replaced by A21 (I24 in SARS-CoV), V49 by I47 (V50), and L160 by V158 (L161). These changes are most likely responsible for a small discrepancy between the ADPr molecules bound to these structures.

**Fig. 6.**
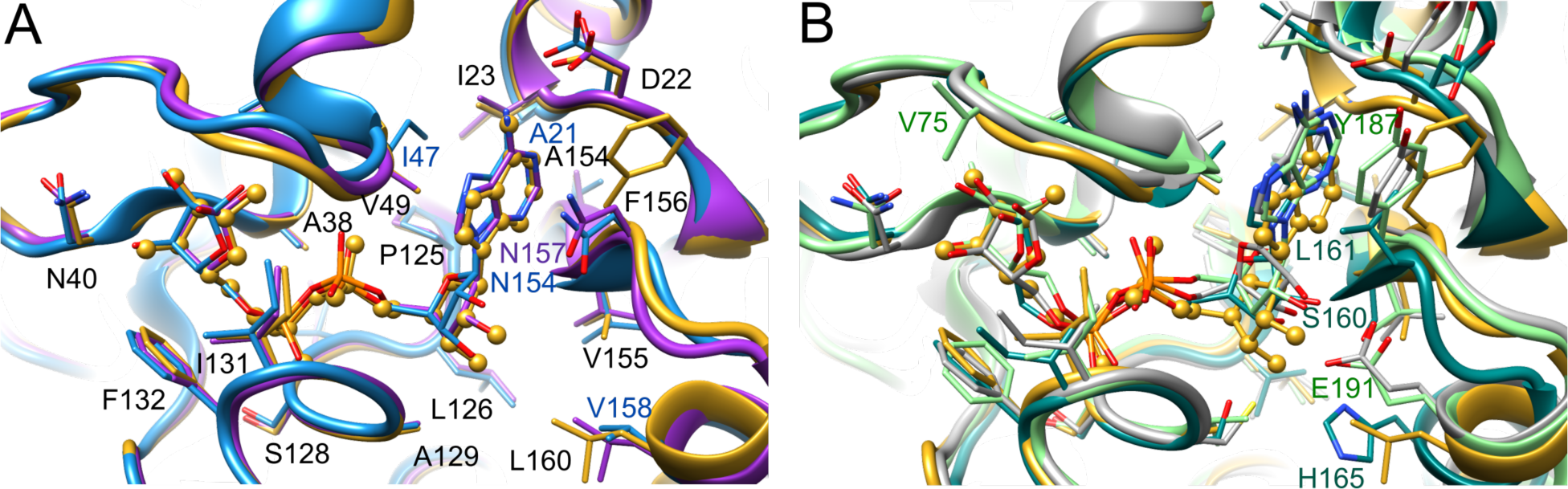
Comparison of SARS-CoV-2 ADRP/ADPr and homologous complexes. **A).** Superposition of SARS-CoV-2 ADRP/ADPr (yellow) with MERS-CoV (5DUS, blue) and SARS-CoV (2FAV, purple). B). Superposition of SARS-CoV-2 ADRP/ADPr with H-CoV-229E (3EWR, grey), IBV (3EWP, teal), FIPV (3JZT, green). In A, SARS-CoV-2 residues are labeled in black. In both panels, selected residues of homologous proteins are labeled.

A more diverged picture is observed in distant homologs from H-CoV-229E, IBV, and FIPV (Fig. 6), mainly in the A and R1 sites, with caveat that the distal ribose in H-CoV-229E ADRP complex has wrong stereochemistry. In these homologs, we observe sequence variation in the F156 position, which is replaced by other hydrophobic residues. The adenine ring is significantly shifted with respect to SARS-CoV-2 ADRP. The interaction between N1 of adenine and D22 equivalent is lost, even though the latter amino acid is conserved in the 3-dimensional context (in IBV, D20 does not overlap in the primary sequence). The distal ribose is better anchored in place – hydrogen bonds link it either to the glutamate residue (E156 in H-CoV-229E, E191 in FIPV) that substitutes L160, or serine in position of V155 (S160 in IBV). Another notable difference is observed in the R2 site, where equivalents of I131 in H-CoV-229E and IBV proteins adopt outlier rotamers, yet the electron density maps allow to model more favorable conformations seen in our structure. In these two models, the proximal ribose adopts α configuration on the anomeric carbon atom (Fig. 6). Such a state, with partial occupancy was also reported for one of the SARS-CoV/ADPr complexes (Egloff *et al.*, 2006) and is linked to the alternative, apo-like conformation of G47-G48. The α configuration most likely illustrates the geometry of the putative substrate, as only then the hydroxyl group is exposed to the solvent, providing room for the macromolecule portion of the substrate.

The common feature in the R2 site among all homologs is the presence of equivalents of F132, Asn40 and glycine-rich loop – these elements through mutational studies have been shown to be crucial for ADRP activity of the SARS-CoV protein (Egloff *et al.*, 2006, Li *et al.*, 2016) and macrodomains from viruses from other families (Malet *et al.*, 2009, Li *et al.*, 2016).

### Catalytic mechanism

In the absence of potential catalytic residues conserved across all the macrodomains, Jankevicius et al. proposed the enzymatic mechanisms involving substrate-assisted catalysis, where a water molecule responsible for nucleophilic attack on the ribose anomeric carbon atom is activated by the Pα group (Jankevicius *et al.*, 2013). In the current ADRP/ADPr structure, the candidate water molecule (Wat) binds to the amide group of A50, carbonyl of A38, oxygen atom of Pα and OH1’ group of the proximal ribose ring of ADPr (Fig. 3, 4). In the ADRP/APO2 and ADRP/MES structures, the last hydrogen bond is replaced by interaction with carbonyl group of G47, enhancing proton abstraction capabilities of the environment. Presumably, based on the models where ADPr exists as α anomer, similar network would be likely to occur in the complex with ADPr-protein or RNA substrates, assuming no major conformational rearrangements occur. The water molecule is ideally located to pursue a nucleophilic attack in the anomeric carbon atom.

## Conclusions

A large, multidomain Nsp3 includes ADP-ribose phosphatase domain (ADRP/MacroD) which is believed to interfere with the host immune response by removing ADP-ribose from ADP-ribosylated proteins or RNA. Our study presents five, atomic and high-resolution structures of SARS-CoV-2 ADRP, including the apo form and complexes with MES, AMP and ADPr. The analysis shows that the enzyme undergoes conformational change upon ADPr binding, which is in agreements with several previous data showing such rearrangements. The shifts affecting both main chain and side chains, are observed primarily around proximal ribose, where the protein has to make room for the sugar moiety and adjust to both configurations of the anomeric carbon atom. The active site water molecule is proposed to carry out nucleophilic attack on the ribose anomeric carbon atom. Our high-resolution studies of ADRP complexes with ligands allow for accurate modeling the active site of ADRP and aid design compounds that can inhibit the activity of this enzyme.

## Materials and methods

### Gene cloning, protein expression and purification

The gene for ADRP was synthesized using codon optimization algorithm for *E. coli* expression and was cloned into pET15b vector (Bio Basic) and transformed into the *E. coli* BL21(DE3)-Gold strain (Stratagene). For preparative purposes, for each protein batch a 4 L culture of LB Lennox medium was grown at 37°C (190 rpm) in presence of ampicillin 150 μg/ml. Once the culture reached OD600 ∼1.0, the temperature setting was changed to 4°C. When bacterial suspension cooled down to 18°C it was supplemented with the following components to indicated concentration: 0.2 mM IPTG, 0.1% glucose, and 40 mM K2HPO4. The temperature was set to 18°C for 20 hours incubation. Bacterial cells were harvested by centrifugation at 7,000 *g* and cell pellets were collected.

We have developed two protocols for purification, differing in buffer composition. For the first batch of the protein (ADRP(b1)) HEPES/NaOH pH 8.0 was used as primary buffering component while subsequent purifications (ADRP(b2)) used Tris/HCl with identical pHs, unless stated otherwise. All the steps were the same, as follows. The cell pellets resuspended in a 12.5 ml lysis buffer (500 mM NaCl, 5% (v/v) glycerol, 50 mM HEPES (or Tris) pH 8.0, 20 mM imidazole, and 10 mM β-mercaptoethanol in Tris-based purification) per liter culture and sonicated at 120W for 5 minutes (4sec ON, 20 sec OFF). The insoluble material was removed by centrifugation at 30,000 *g* for one hour at 4°C. Supernatant was mixed with 3 ml of Ni^2+^ Sepharose (GE Healthcare Life Sciences) equilibrated with lysis buffer with imidazole concertation brought up to 50 mM and suspension was applied on Flex-Column (420400-2510) connected to Vac-Man vacuum manifold (Promega). Unbound protein was washed out through controlled suction with 160 ml of lysis buffer (50 mM imidazole). Bound protein was eluted with 15 ml of lysis buffer supplemented to 500 mM imidazole pH 8.0. 2 mM DTT was added and the protein was subsequently treated overnight at 4°C with Tobacco Etch Mosaic Virus (TEV) protease at 1:20 protease:protein ratio. The protein solution was concentrated on 10 kDa MWCO filter (Amicon-Millipore) and further purified on size exclusion column Superdex 200 in lysis buffer where β-mercaptoethanol was replaced with 1 mM TCEP. Fraction containing ADRP were pooled and run one more time through Ni^2+^ Sepharose. Flow through was collected and buffer-exchanged to crystallization buffer (150 mM NaCl, 20 mM HEPES pH 7.5 (or Tris pH 8.0), 1 mM TCEP) via 10X concentration/dilution repeated 3 times. The protein was immediately used for crystallization trials. Final concentrations of ADRP(b1) was 22 mg/ml. Final concentrations of ADRP(b2) was 32 mg/ml.

### Crystallization

Crystallization screens were performed by the sitting-drop vapor-diffusion method in 96-well CrystalQuick plates (Greiner Bio-One). The plates were set up using Mosquito liquid dispenser (TTP LabTech) utilizing 400 nl of purified protein sample was mixed with 400 nl of well solution and equilibrated against 135 nl reservoir solution. ADRP(b1) was used to grow apo from crystals and in crystallization with AMP and ADPr. The AMP complexes was prepared by adding AMP (pH 6.5) to a final concertation of 12 mM. To obtain ADPr complex the protein was mixed with ADPr in the molar proportion of 1:2. The screening was performed using MCSG1, MCSG4 (Anatrace), SaltRX (Hampton), PACT (Qiagen), and INDEX (Hampton) screens. ADRP(b2) was set up at 18 mg/mL with Pi-minimal (Jena Biosciences), Protein Complex (NeXtal), and INDEX (Hampton) screens. In all cases, the plates were incubated at 289 K.

ADRP(b1) crystals grew from conditions containing 0.1 M CHES pH 9.5, 30% (w/v) PEG3000, yielding a structure denoted ADRP/APO1. Complex with ADPr was obtained from 0.01 M sodium citrate, 33% PEG6000, giving the structure labeled as ADRP/ADPr. Complex with AMP was grown from 0.1 M MES pH 6.5, 30% (w/v) PEG4000, giving structures labeled as ADRP/AMP. ADRP(b2) crystals grew from 0.1 M MES, pH 6.5, 30% (w/v) PEG 4000, yielding ADRP/MES complex, and from 30 mM Na/K tartrate, 150 mM AMPD/Tris pH 9.0, 34.3 % (w/v) PEG5000 MME, giving the ADRP/APO2 crystals.

### Data collection, structure determination and refinement

Prior to flash-cooling in liquid nitrogen, the crystals were cryoprotected in their mother liquor supplemented with: the increased concentration of PEG3000 up to 40% (ADRP/APO1), 5% glycerol (ADRP/ADPr), 7% (ADRP/AMP) or 10% ethylene glycol (ADRP/MES). The ADRP/APO2 crystals did not require cryoprotection. The X-ray diffraction experiments were carried out at 100 K, at the Structural Biology Center 19-ID beamline at the Advanced Photon Source, Argonne National Laboratory. The diffraction images were recorded on the PILATUS3 X 6M detector. The data set was processed and scaled with the HKL3000 suite (Minor *et al.*, 2006). Intensities were converted to structure factor amplitudes in the truncate program (French & Wilson, 1978, Padilla & Yeates, 2003) from the CCP4 package (Winn *et al.*, 2011). The ADRP/APO1 structure was determined by molecular replacement (MR) using Molrep (Vagin & Teplyakov, 2010) implemented in the HKL3000 software package and SARS-CoV ADRP structure (PDB id 2ACF) as a search probe. The subsequent structures where solved by MR with a refined SARS-CoV-2 ADRP as a model. In all cases, the initial solution was manually adjusted using COOT (Emsley & Cowtan, 2004) and then iteratively refined using Coot, Phenix (Adams *et al.*, 2010) and Refmac (Murshudov *et al.*, 1997, Winn *et al.*, 2011). The final rounds of refinement were carried out in Phenix (ADRP/APO1, ADRP/ADPr, ADRP/MES) or Refmac (ADRP/AMP, ADRP/APO2). ADRP/APO1 and ADRP/ADPr were refined with TLS parameterization of anisotropic displacement parameters while for the remaining structures full anisotropic refinement was calculated. Throughout the refinement, the same 5% of reflections were kept out throughout from the refinement (in both REFMAC and PHENIX refinements). The final models show nearly complete polypeptide chains. The residues that have not been modeled due to the lack of interpretable electron density include: Gly1-Glu2 and Glu170 in chains A and B for ADRP/APO1, Gly1-Glu2-Val3, Leu169-Glu170 in chain A and Gly1-Glu2, Glu170 in chain B for ADRP/ADPr, Gly1-Glu2 in chain A and Gly1-Glu2, Glu170 in chain B for ADRP/AMP, Gly1-Glu2-Val3 and Glu170 for ADRP/MES, and Gly1-Glu2 for ADRP/APO2. The stereochemistry of the structure was checked with MolProbity (Davis *et al.*, 2007) PROCHECK (Laskowski *et al.*, 1993) and the Ramachandran plot and validated with the PDB validation server. The data collection and processing statistics are given in Table 1. The atomic coordinates and structure factors have been deposited in the PDB under accession codes 6VXS, 6W02, 6W6Y, 6WCF, and 6WEN.

## Acknowledgements

We truthfully thank the members of the SBC at Argonne National Laboratory, especially Darren Sherrell and Alex Lavens for their help with setting beamline and data collection at beamline 19-ID, and Paula Bulaon for help with manuscript editing. Funding for this project was provided in part by federal funds from the National Institute of Allergy and Infectious Diseases, National Institutes of Health, Department of Health and Human Services, under Contract No. HHSN272201700060C. The use of SBC beamlines at the Advanced Photon Source is supported by the U.S. Department of Energy (DOE) Office of Science and operated for the DOE Office of Science by Argonne National Laboratory under Contract No. DE-AC02-06CH11357.

## References

Adams, P. D., Afonine, P. V., Bunkoczi, G., Chen, V. B., Davis, I. W., Echols, N., Headd, J. J., Hung, L. W., Kapral, G. J., Grosse-Kunstleve, R. W., McCoy, A. J., Moriarty, N. W., Oeffner, R., Read, R. J., Richardson, D. C., Richardson, J. S., Terwilliger, T. C. & Zwart, P. H. (2010). Acta Crystallogr., Sect D: Biol. Crystallogr. 66, 213–221.

Atasheva, S., Akhrymuk, M., Frolova, E. I. & Frolov, I. (2012). J Virol 86, 8147–8160.

Atasheva, S., Frolova, E. I. & Frolov, I. (2014). J Virol 88, 2116–2130.

Baez-Santos, Y. M., St John, S. E. & Mesecar, A. D. (2015). Antiviral Res. 115, 21–38.

Cho, C. C., Lin, M. H., Chuang, C. Y. & Hsu, C. H. (2016). J Biol Chem 291, 4894–4902.

Coronaviridae Study Group of the International Committee on Taxonomy of Viruses (2020). Nat Microbiol 5, 536–544.

Crawford, K., Bonfiglio, J. J., Mikoc, A., Matic, I. & Ahel, I. (2018). Crit Rev Biochem Mol Biol 53, 64–82.

Cui, J., Li, F. & Shi, Z. L. (2019). Nat. Rev. Microbiol. 17, 181–192.

Davis, I. W., Leaver-Fay, A., Chen, V. B., Block, J. N., Kapral, G. J., Wang, X., Murray, L. W., Arendall, W. B., 3rd, Snoeyink, J., Richardson, J. S. & Richardson, D. C. (2007). Nucleic Acids Res. 35, W375–383.

Davis, I. W., Murray, L. W., Richardson, J. S. & Richardson, D. C. (2004). Nucleic Acids Res. 32, W615–619.

Eckei, L., Krieg, S., Butepage, M., Lehmann, A., Gross, A., Lippok, B., Grimm, A. R., Kummerer, B. M., Rossetti, G., Luscher, B. & Verheugd, P. (2017). Sci Rep 7, 41746.

Egloff, M. P., Malet, H., Putics, A., Heinonen, M., Dutartre, H., Frangeul, A., Gruez, A., Campanacci, V., Cambillau, C., Ziebuhr, J., Ahola, T. & Canard, B. (2006). J Virol 80, 8493–8502.

Emsley, P. & Cowtan, K. (2004). Acta Crystallogr., Sect D: Biol. Crystallogr. 60, 2126–2132.

Fontana, P., Bonfiglio, J. J., Palazzo, L., Bartlett, E., Matic, I. & Ahel, I. (2017). Elife 6.

French, S. & Wilson, K. (1978). Acta Crystallogr., Sect. A: Found. Crystallogr. 34, 517–525.

Grunewald, M. E., Chen, Y., Kuny, C., Maejima, T., Lease, R., Ferraris, D., Aikawa, M., Sullivan, C. S., Perlman, S. & Fehr, A. R. (2019). PLoS Pathog 15, e1007756.

Hintze, B. J., Lewis, S. M., Richardson, J. S. & Richardson, D. C. (2016). Proteins 84, 1177–1189.

Holm, L. & Rosenstrom, P. (2010). Nucleic Acids Res. 38 Suppl, W545–549.

Jankevicius, G., Hassler, M., Golia, B., Rybin, V., Zacharias, M., Timinszky, G. & Ladurner, A. G. (2013). Nat Struct Mol Biol 20, 508–514.

Karplus, P. A. & Diederichs, K. (2012). Science 336, 1030–1033.

Kuri, T., Eriksson, K. K., Putics, A., Zust, R., Snijder, E. J., Davidson, A. D., Siddell, S. G., Thiel, V., Ziebuhr, J. & Weber, F. (2011). J Gen Virol 92, 1899–1905.

Laskowski, R. A., MacArthur, M. W., Moss, D. S. & Thornton, J. M. (1993). Journal of Applied Crystallography 26, 283–291.

Lee, H. J., Shieh, C. K., Gorbalenya, A. E., Koonin, E. V., La Monica, N., Tuler, J., Bagdzhadzhyan, A. & Lai, M. M. (1991). Virology 180, 567–582.

Lei, J. & Hilgenfeld, R. (2016). Virol Sin 31, 288–299.

Lei, J., Kusov, Y. & Hilgenfeld, R. (2018). Antiviral Res 149, 58–74.

Li, C., Debing, Y., Jankevicius, G., Neyts, J., Ahel, I., Coutard, B. & Canard, B. (2016). J Virol 90, 8478–8486.

Malet, H., Coutard, B., Jamal, S., Dutartre, H., Papageorgiou, N., Neuvonen, M., Ahola, T., Forrester, N., Gould, E. A., Lafitte, D., Ferron, F., Lescar, J., Gorbalenya, A. E., de Lamballerie, X. & Canard, B. (2009). J Virol 83, 6534–6545.

Minor, W., Cymborowski, M., Otwinowski, Z. & Chruszcz, M. (2006). Acta Crystallogr., Sect D: Biol. Crystallogr. 62, 859–866.

Moss, J., Tsai, S. C., Adamik, R., Chen, H. C. & Stanley, S. J. (1988). Biochemistry 27, 5819–5823.

Munnur, D., Bartlett, E., Mikolcevic, P., Kirby, I. T., Matthias Rack, J. G., Mikoc, A., Cohen, M. S. & Ahel, I. (2019). Nucleic Acids Res 47, 5658–5669.

Murshudov, G. N., Vagin, A. A. & Dodson, E. J. (1997). Acta Crystallogr., Sect D: Biol. Crystallogr. 53, 240–255.

Padilla, J. E. & Yeates, T. O. (2003). Acta Crystallogr., Sect D: Biol. Crystallogr. 59, 1124–1130.

Pehrson, J. R. & Fried, V. A. (1992). Science 257, 1398–1400.

Piotrowski, Y., Hansen, G., Boomaars-van der Zanden, A. L., Snijder, E. J., Gorbalenya, A. E. & Hilgenfeld, R. (2009). Protein science: a publication of the Protein Society 18, 6–16.

Rack, J. G., Perina, D. & Ahel, I. (2016). Annu Rev Biochem 85, 431–454.

Rice, P., Longden, I. & Bleasby, A. (2000). Trends Genet. 16, 276–277.

Saikatendu, K. S., Joseph, J. S., Subramanian, V., Clayton, T., Griffith, M., Moy, K., Velasquez, J., Neuman, B. W., Buchmeier, M. J., Stevens, R. C. & Kuhn, P. (2005). Structure 13, 1665–1675.

Vagin, A. & Teplyakov, A. (2010). Acta Crystallographica Section D-Biological Crystallography 66, 22–25.

Winn, M. D., Ballard, C. C., Cowtan, K. D., Dodson, E. J., Emsley, P., Evans, P. R., Keegan, R. M., Krissinel, E. B., Leslie, A. G., McCoy, A., McNicholas, S. J., Murshudov, G. N., Pannu, N. S., Potterton, E. A., Powell, H. R., Read, R. J., Vagin, A. & Wilson, K. S. (2011). Acta Crystallogr., Sect D: Biol. Crystallogr. 67, 235–242.

Wojdyla, J. A., Manolaridis, I., Snijder, E. J., Gorbalenya, A. E., Coutard, B., Piotrowski, Y., Hilgenfeld, R. & Tucker, P. A. (2009). Acta Crystallogr D Biol Crystallogr 65, 1292–1300.

Wu, F., Zhao, S., Yu, B., Chen, Y. M., Wang, W., Song, Z. G., Hu, Y., Tao, Z. W., Tian, J. H., Pei, Y. Y., Yuan, M. L., Zhang, Y. L., Dai, F. H., Liu, Y., Wang, Q. M., Zheng, J. J., Xu, L., Holmes, E. C. & Zhang, Y. Z. (2020). Nature 579, 265–269.

Xu, Y., Cong, L., Chen, C., Wei, L., Zhao, Q., Xu, X., Ma, Y., Bartlam, M. & Rao, Z. (2009). J Virol 83, 1083–1092.

